# Stress Resets Transgenerational Small RNA Inheritance

**DOI:** 10.1101/669051

**Authors:** Leah Houri-Ze’evi, Guy Teichman, Hila Gingold, Oded Rechavi

## Abstract

Transgenerational inheritance of small RNAs is challenging basic concepts of heredity and achieving control over such responses is of great interest. In *C. elegans* nematodes, small RNAs are transmitted across generations to establish a transgenerational memory trace of ancestral environments and distinguish self from non-self genes. Inheritance of small RNAs is regulated by dedicated machinery and carryover of aberrant heritable small RNA responses was shown to be maladaptive and to induce sterility. Here we show that various types of stress (starvation, high temperatures, and high osmolarity) but not non-stressful changes in cultivation conditions, lead to resetting of small RNA inheritance. We found that stress leads to a genome-wide reduction in heritable small RNA levels and that mutants defective in different stress pathways exhibit irregular RNAi inheritance dynamics. Moreover, we discovered that resetting of heritable RNAi is orchestrated by MAPK pathway factors, the transcription factor SKN-1, and the MET-2 methyltransferase. Termination of small RNA inheritance, and the fact that this process depends on stress, could protect from run-on of environment-irrelevant heritable gene regulation.

## Introduction

Different human diseases, for example a number of imprinting-associated syndromes (Angelman syndrome, Prader-Willi syndrome, and Beckwith-Wiedemann syndrome) arise due to inheritance of parental information which is not encoded in the DNA sequence (Tucci et al., 2019). In addition, while the mechanisms are still unclear, many widespread disorders were suggested to be influenced by non-genetic inheritance and to be affected by the ancestors’ life history (Bohacek and Mansuy, 2015; Chen et al., 2016; Gapp et al., 2014; Kazachenka et al., 2018; Nilsson et al., 2018; Öst et al., 2014; Skvortsova et al., 2018; Teperino et al., 2013). Thus, development of methods for resetting of detrimental heritable effects could potentially benefit future treatment of these diseases and enable the progeny to start as a “blank slate”.

In *Caenorhabditis elegans* nematodes, much knowledge has been gained in recent years regarding the mechanisms that enable transgenerational gene regulation via inheritance of small RNAs (Rechavi and Lev, 2017; Serobyan and Sommer, 2017). Dedicated small RNA inheritance proteins, which are required specifically for heritable responses, have been identified (de Albuquerque et al., 2015; Ashe et al., 2012; Buckley et al., 2012; Houri-Ze’evi et al., 2016; Lev et al., 2017, 2019; Shirayama et al., 2012; Wan et al., 2018; Xu et al., 2018)). In worms, multigenerational small RNA inheritance is robust and requires RNA-dependent RNA Polymerases (RdRPs) which use the target mRNA as a template for amplifying “Secondary” and “Tertiary” small RNAs (Rechavi et al., 2011; Sapetschnig et al., 2015). The amplification reaction outcompetes the dilution of the heritable small RNA molecules in every generation (Rechavi et al., 2011). Multiple “Primary” small RNA species (exogenous and endogenous small interfering RNAs and PIWI-interacting RNAs) can induce amplification of heritable small RNAs (Das et al., 2008; Sijen et al., 2001). In the nucleus, amplified small RNAs lead to transcriptional silencing of their targets, in cooperation with chromatin regulators (Burton et al., 2011; She et al., 2009). Nuclear small RNAs modify the chromatin, and some changes in histone marks are transgenerationally inherited, also in response to environmental changes (Klosin et al., 2017). Specific worm Argonaute proteins (HRDE-1, CSR-1, WAGO-4, WAGO-1) were found to physically carry small RNAs in the germline and across generations (Ashe et al., 2012; Claycomb et al., 2009; Shirayama et al., 2012; Wedeles et al., 2013; Xu et al., 2018)

The worm’s small RNA pools change transgenerationally in response to multiple environmental challenges such as viral infection (Rechavi et al., 2011), starvation (Rechavi et al., 2014), and stressful temperatures (Ni et al., 2016; Schott et al., 2014). Further, *C. elegans* actively regulates small RNA inheritance and controls the duration and potency of the transgenerational effects (Houri-Zeevi and Rechavi, 2017). Heritable RNA interference (RNAi) responses, which are mediated by small RNAs, can be induced by targeting of germline-expressed genes using double-stranded RNA (dsRNA) triggers. Typically, such heritable responses last 3-5 generations (Alcazar et al., 2008), but the duration of the heritable response can differ in some mutants. For example, MET-2, an histone 3 lysine 9 (H3K9) methyltransferase, is required for termination of heritable RNAi, and accordingly RNAi inheritance is stable in *met-2* mutants (Lev et al., 2017). Moreover, even in wild type animals, inheritance of ancestral RNAi responses can be extended by triggering dsRNA-induced silencing of other genes in the progeny (Houri-Ze’evi et al., 2016). Together, these different mechanisms constitute a transgenerational “timer” which restricts inheritance of small RNA responses (Houri-Zeevi and Rechavi, 2017).

It was hypothesized that continuation of some gene expression programs in the progeny could increase the descendants’ chances to survive, especially if parents and progeny experience the same conditions (Houri-Ze’evi et al., 2016; Jablonka, 2013, 2017; Kishimoto et al., 2017). However, if the environmental conditions change, the carryover of ancestral responses could become detrimental (Jablonka, 2013). Indeed, in worms, mutants which are unable to regulate multigenerational accumulation of heritable small RNAs become sterile (Lev et al., 2017, 2019; Ni et al., 2016; Simon et al., 2014). Therefore, it is possible that mechanisms have evolved to terminate or extend small RNA-based inheritance according to the presence or absence of dramatic shifts in settings.

In this study, we examined if changes in environmental conditions between generations alter the dynamics of small RNA inheritance. We found that stress resets small RNA inheritance and that termination is orchestrated by the MAPK stress pathway, the SKN-1 transcription factor, and the MET-2 H3K9 methyltransferase. We suggest that this mechanism could unburden the progeny from irrelevant heritable gene regulation.

## Results

We examined whether discrepancies between the growth conditions of the ancestors and their progeny would affect small RNA inheritance. We applied different types of stress to progeny that inherited small RNA responses from unstressed parents. To monitor small RNA-mediated inheritance, we used three different inheritance assays:

First, we investigated the heritable responses initiated by exogenously-derived small RNAs, by silencing a single-copy germline-expressed *gfp*. Feeding the worms with bacteria that express anti-*gfp* directed dsRNA induces heritable silencing that lasts for ±3-5 consecutive generations (Alcazar et al., 2008) (**Methods**). We subjected P0 animals to RNAi and tested how the heritable silencing is affected in the F1 progeny following exposure to three different types of stressors, experienced during the first larval stage (L1). To stress the worms we heat shocked the progeny for 2 hours (Zevian and Yanowitz, 2014), cultured the progeny in hyperosmotic conditions for 48 hours (Rodriguez et al., 2013), or starved the progeny for 6 days (Rechavi et al., 2014) (See **Figure 1A, 1B** and **Methods**). We then examined the inheritance of RNAi responses that were initiated in the previous non-stressed generation. We found that stressing the worms during the L1 stage led to a reduction in heritable GFP silencing at the adult stage. Importantly, the strong reduction in silencing was not restricted to the F1 progeny that were directly stressed as it was observed also in the non-stressed F2 and F3 progeny (q-value<0.0004, **Figure 1C**). Thus, stress experienced during the F1 generation (at the L1 stage) erases dsRNA-induced (exo-siRNA) silencing in a transgenerational manner.

**Figure 1:**
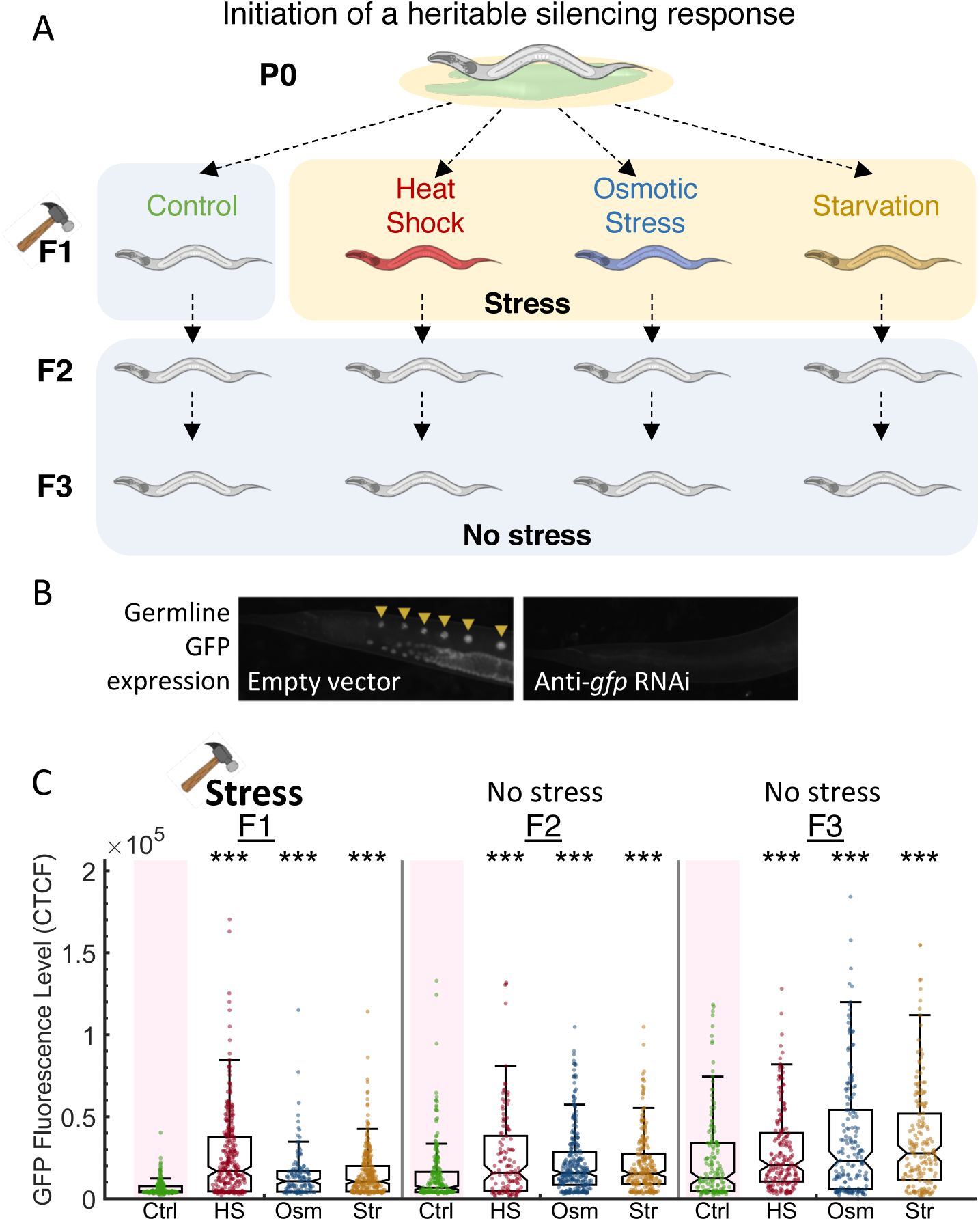
Stress resets heritable small RNA silencing. A. *Experimental scheme*. Heritable small RNA responses are initiated at the first generation, and the F1 progeny are then subjected to three different stress types (heat shock {HS}, hyperosmotic stress {Osm}, starvation {Str}). Inheritance of the ancestral response is scored both in the stressed generation and in the next generations which were grown under regular growth conditions. B. Representative images of worms containing the *Pmex-5::gfp* transgene, treated with empty vector containing bacteria (left) or with anti-*gfp* dsRNA-producing bacteria (right). C. *Heritable exo-siRNAs silencing is reset by stress.* The graph displays the measured germline GFP fluorescence levels of individual worms (y-axis) across generations under the indicated condition (x-axis). Each dot represents the value of an individual worm. Shown are the median of each group, with box limits representing the 25^th^ (Q1) and 75^th^ (Q3) percentiles, notch representing a 95% confidence interval, and whiskers indicating Q1-1.5*IQR and Q3+1.5*IQR. FDR-corrected values were obtained using Dunn’s test. (***) indicates q<0.001 (see **Methods**).

Next, we examined if stress also resets heritable silencing that is triggered by endogenous small interfering RNAs (endo-siRNAs) or PIWI-interacting small RNAs (piRNAs). For this purpose, we used (1) worms that carry a *gfp* transgene that contains an endo-siRNA-target sequence (an “endo-siRNA” sensor, integrated in the genome of the GR1720 strain (Billi et al., 2012)), and (2) worms that stochastically silence a foreign *mcherry* transgene, that contains multiple piRNAs-recognition sites (Zhang et al., 2018) (see also **Figure 2A and 2B**, and **Methods)**. We found that all three stressors (starvation, high temperatures, and hyperosmotic conditions) reset both endo-siRNAs- and piRNAs-mediated heritable silencing in worms directly exposed to stress (q-value<0.0005 and q-value<0.0007, **Figure 2A** and **2B**). However, in contrast to exo-siRNAs mediated silencing that was reset in a transgenerational manner (**Figure 1C**), we found that unstressed worms in the next generations re-established endo-siRNAs- and piRNAs-mediated silencing. Unlike exogenous dsRNA-derived primary siRNAs which cannot be re-synthesized in the progeny, primary piRNAs and primary endo-siRNAs are encoded in the genome and therefore we reason that these small RNAs are transcribed *de novo* in each generation, compensating in the next generation for stress-induced erasure of the parental molecules.

**Figure 2:**
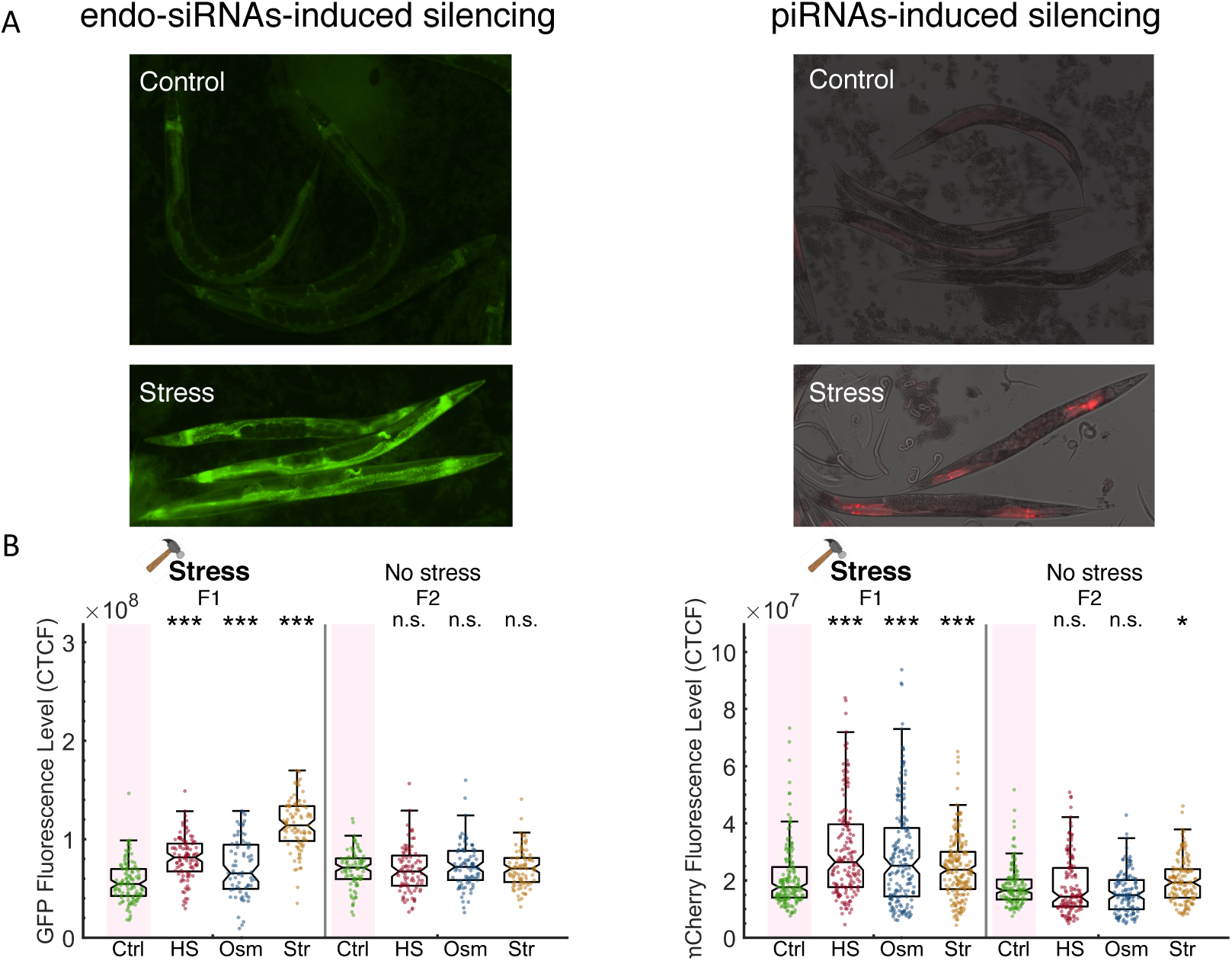
Stress resets endo-siRNAs and piRNAs-induced silencing. A. Representative images of worms expressing endo-siRNAs sensor (left) or the piRNAs sensor (right) under control or stress (HS) conditions. B. The graph displays the measured GFP (left, endo-siRNAs sensor) or mCherry (right, piRNAs sensor) fluorescence levels of individual worms (y-axis) across generations under the indicated condition (x-axis). Each dot represents the value of an individual worm. Shown are the median of each group, with box limits representing the 25^th^ (Q1) and 75^th^ (Q3) percentiles, notch representing a 95% confidence interval, and whiskers indicating Q1-1.5*IQR and Q3+1.5*IQR. FDR-corrected values were obtained using Dunn’s test. Not significant (ns) indicates q ≥ 0.05, (*) indicates q<0.05, and (***) indicates q<0.001 (see **Methods**).

We then tested whether the ability of stress to reset heritable small RNA responses depends on the developmental stage or the generation in which stress is applied after the initial RNAi event:

As was observed when stress was applied during the L1 stage, we found that stressing adult worms also led to resetting of heritable silencing (q<0.0001, **Supplementary Figure 1**). Thus, stress-induced resetting of RNAi inheritance is not restricted to the L1 stage.

While exposing the F1 generation to stress led to erasure of the heritable silencing response (as described in **Figure 1B**), when stress was applied two generations after exposure to dsRNA, we did not detect statistically-significant stress-induced resetting (**Figure 3A**). In accordance with these results, we previously showed that re-challenging the F1 but not F2 progeny with RNAi extends the duration of ancestral RNAi responses (Houri-Ze’evi et al., 2016). Together, these results reveal a critical period, at the F1 generation, during which heritable silencing is still plastic.

**Figure 3:**
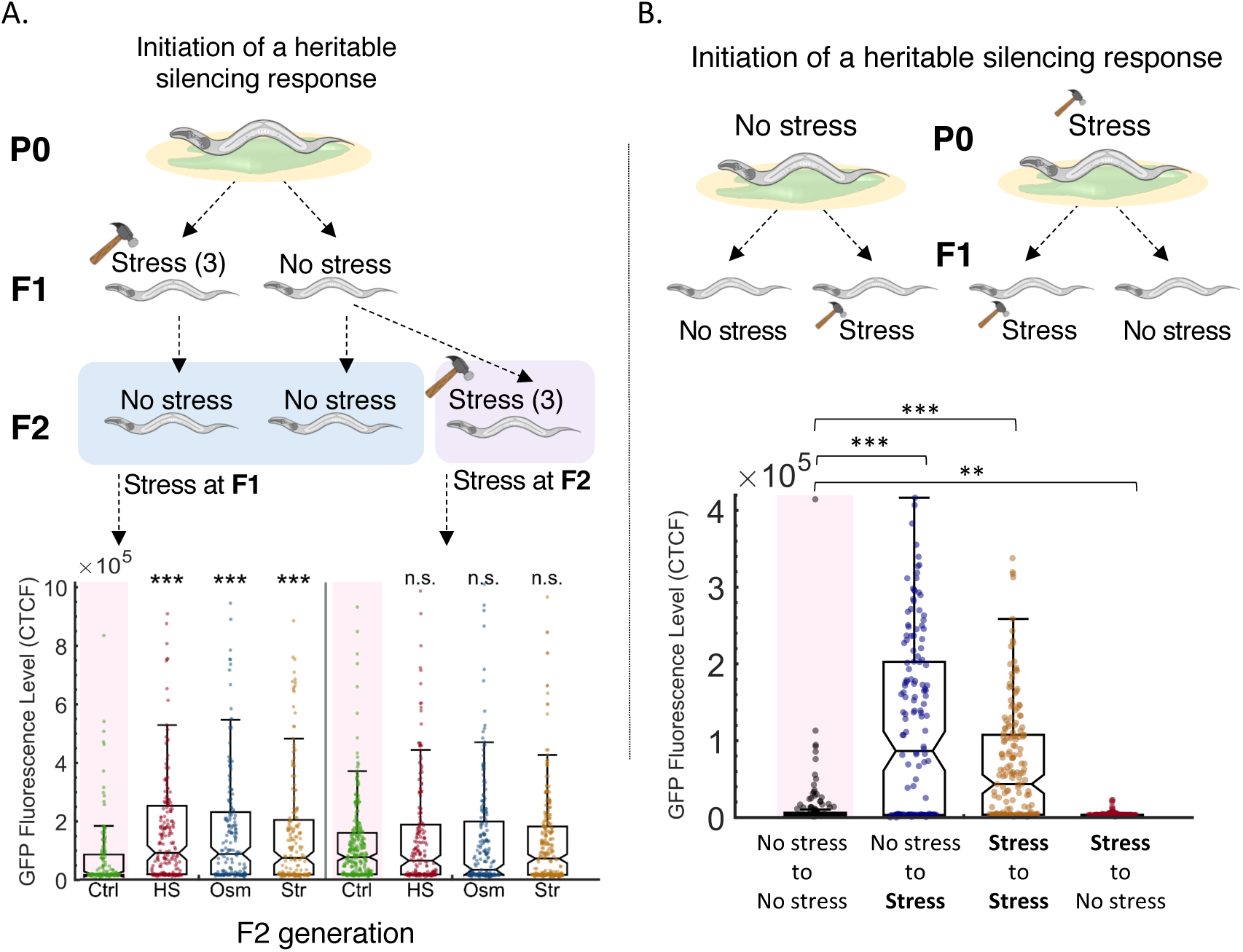
Resetting of heritable small RNA responses can only occur during the F1 generations and is induced specifically by stress. A. Stress applied two generations after the initiation of the heritable small RNAs response does not reset the inheritance. Upper panel: *Experimental scheme*. A heritable anti-*gfp* small RNA response is initiated at the first generation and the progeny are then subjected to three different stress types (heat shock, hyperosmotic stress, starvation). Stress is applied at the first generation (F1) or the second generation (F2) after the initiation of the heritable response. Inheritance of the ancestral response is scored at the F2 generation. Lower panel: *The ability of stress to reset heritable small RNAs depends on the generation during which stress is applied.* Worms which were stressed two generations after the initiation of RNAi did not show any altered inheritance dynamics. The graph displays the measured GFP fluorescence levels of individual worms (y-axis) across generations under the indicated condition (x-axis). Each dot represents the value of an individual worm. Shown are the median of each group, with box limits representing the 25^th^ (Q1) and 75^th^ (Q3) percentiles, notch representing a 95% confidence interval, and whiskers indicating Q1-1.5*IQR and Q3+1.5*IQR. FDR-corrected values were obtained using Dunn’s test. Not significant (ns) indicates q ≥ 0.05 and (***) indicates q<0.001 (see **Methods**). B. Shifting from stress to non-stress conditions fails to reset small RNA inheritance. Upper panel: *Experimental scheme*. Worms grown in regular growth conditions (20°C, control) or in high temperatures (25°C, stress) are exposed to an anti-*gfp* RNAi trigger. The next generation are then grown either in similar conditions (control to control, stress to stress) or transferred to the other growth condition (control to stress, stress to control). Lower panel: *Stress, and not any change in the environment, resets heritable small RNAs*. Worms which were exposed to high temperatures at the next generation reset the RNAi-induced *gfp* silencing regardless of the growth conditions at the previous generation. The graph displays the measured GFP fluorescence levels of individual worms (y-axis) under the indicated condition (x-axis). Each dot represents the value of an individual worm. Shown are the median of each group, with box limits representing the 25^th^ (Q1) and 75^th^ (Q3) percentiles, notch representing a 95% confidence interval, and whiskers indicating Q1-1.5*IQR and Q3+1.5*IQR. FDR-corrected values were obtained using Dunn’s test. not significant (ns) indicates q ≥ 0.05, (**) indicates q<0.01 and (***) indicate q<0.001 (see **Methods**).

Exposure to stress can be viewed simply as a change in the growing conditions, from non-stressful to stressful settings. We wondered whether any form of change – and not just stress – could lead to resetting of heritable small RNA responses. To test this hypothesis, we examined how an “improvement” in the cultivation conditions would affect small RNA inheritance. We exposed parents that were grown under stressful conditions to RNAi and examined the heritable response in progeny grown in non-stressful conditions (see scheme in **Figure 3B**). We found that resetting occurs only in response to stress (q-value<0.0001, **Figure 3B**), and not when the progeny of stressed worms is transferred to non-stressful conditions. In contrast, we observed that RNAi responses initiated in stressed parents are strengthened when the progeny is transferred to non-stressful conditions (q-value<0.0053, **Figure 3B**). Accordingly, RNAi inheritance was weaker in worms that were grown in stressful conditions for two consecutive generations than in worms maintained in non-stressful conditions for the same time (q-value<0.0001, **Figure 3B**). Thus, stress, and not just any change, leads to resetting of heritable small RNA responses, regardless of the ancestral growth conditions.

To study whether stress-induced resetting is a regulated process, we examined mutants defective in different stress pathways. Out of the 9 mutants defective in stress responses that we examined, only two (*daf-16, aak-1/2*) did not show altered RNAi inheritance dynamics. Interestingly, *daf-16* was shown to be involved in heritable “mortal germline” phenotype (Mrt) (Simon et al., 2018) and *aak-1/2* was implicated in regulating transgenerational consequences of starvation (Demoinet et al., 2017). The remaining 7 mutants showed either defective (*mek-1/sek-1, kgb-1, hsf-1*) or enhanced (*pmk-1, sek-1, skn-1, daf-2*) RNAi inheritance (**Figure 4A** and **Supplementary Figure 2** and **3**). This was true irrespectively of whether the mutant progeny experienced stress or not and could suggest that stress regulation and small RNA inheritance are interconnected.

**Figure 4:**
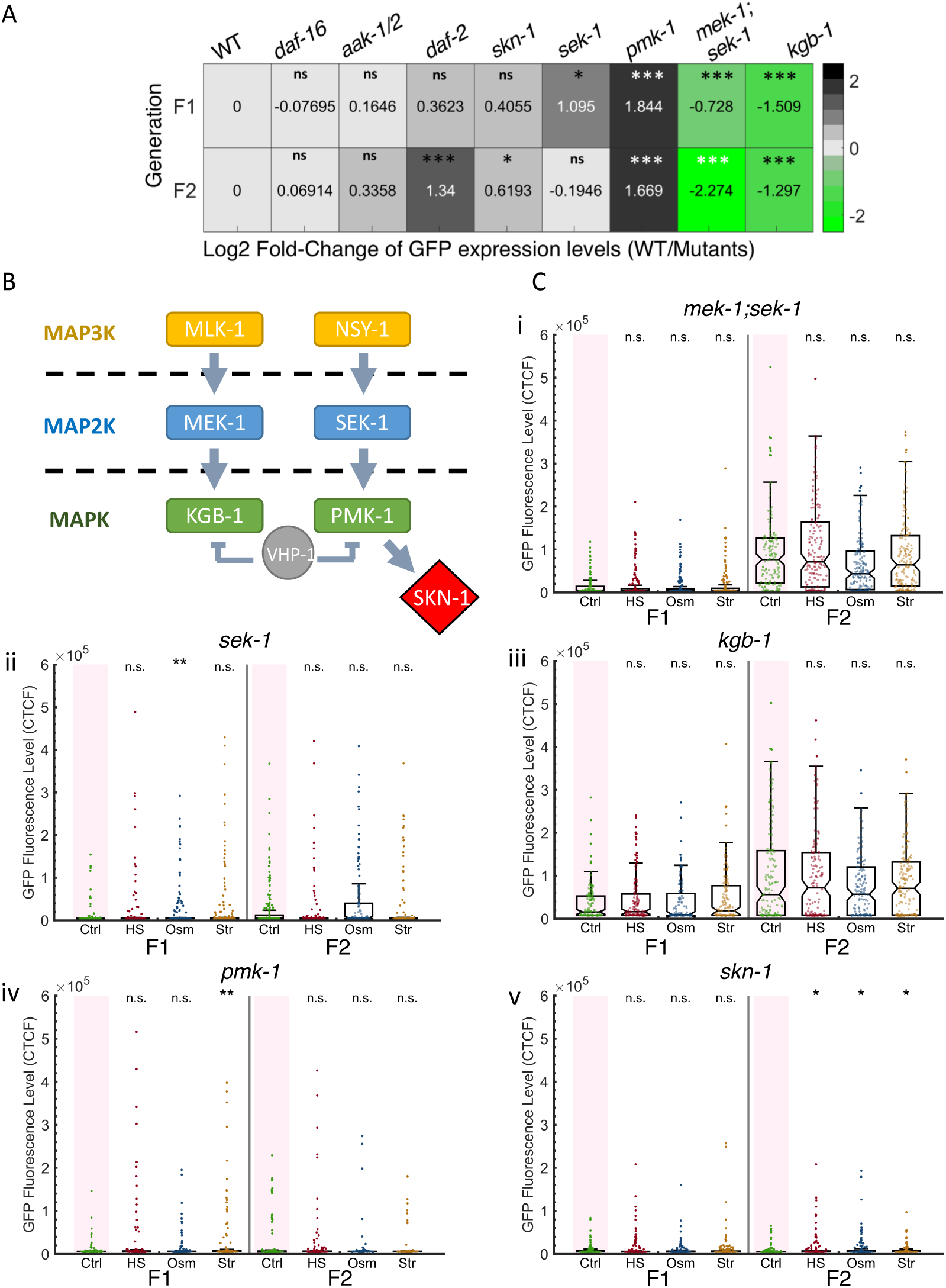
The MAPK pathway and the SKN-1 transcription factor regulate small RNAs resetting in response to stress. A. *Mutants defected in stress processing pathways display altered heritable RNAi dynamics*. Heatmap representing the log_2_-fold change of GFP fluorescence levels (color coded and indicated numbers) in mutants worms compared to WT at the F1 (up) and F2 (down) generations after RNAi. Shown are results under normal conditions (no stress). FDR-corrected values were obtained using Dunn’s test. The comparison values of each mutant were calculated based on its independent experiments (see full results in **Supplementary Figure 2** and **3**). B. Members of *C. elegans’* MAPK pathway, found to influence resetting of small RNAs. Based on (Ewbank, 2006; Inoue et al., 2005; Kim et al., 2004; Mizuno et al., 2008) C. *Mutants defected in the MAPK signaling pathway do not reset heritable RNAi responses following stress*. The graphs display the measured germline GFP fluorescence levels of mutant worms (y-axis) across generations under the indicated condition (x-axis). Each dot represents the value of an individual worm. Each mutant was examined in an independent experiment. Shown are the median of each group, with box limits representing the 25^th^ (Q1) and 75^th^ (Q3) percentiles, notch representing a 95% confidence interval, and whiskers indicating Q1-1.5*IQR and Q3+1.5*IQR. FDR-corrected values were obtained using Dunn’s test. Not significant (ns) indicates q ≥ 0.05, (*) indicates q<0.05, (**) indicates q<0.01, and (***) indicates q<0.001, see **Methods**). Full results (including the side-by-side wild-type results of each experiment) can be found in **Supplementary Figure 2** and **3**.

We next examined whether these genes are not only involved in RNAi inheritance but also in its resetting in response to stress. We found that MAP Kinase (MAPK) genes (*sek-1/mek-1, sek-1, pmk-1, kgb-1*), and the *skn-1* gene (encoding a transcription factor which is regulated by p38 MAPK-dependent phosphorylation) are required specifically for resetting of RNAi inheritance in response to stress (**Figure 4B, 4C**, and **Supplementary Figure 2** and **3**).

The MAPK pathway is necessary for integrating responses to multiple types of stress, such as DNA damage (Bianco and Schumacher, 2018; Ermolaeva et al., 2013), osmotic stress (Gerke et al., 2014), heat shock (Mertenskötter et al., 2013), and pathogen infection (Troemel et al., 2006). Interestingly, we found that some of the mutants that did not reset heritable silencing following stress showed enhanced RNAi inheritance (*sek-1*, *pmk-1, skn-1*, acting in the same signaling cascade), while others were defective in RNAi inheritance (*sek-1/mek-1* and kgb*-1*) (**Figure 4**). Thus, the ability to reset heritable responses following stress, and the ability to transmit RNAi, appear to be two distinct functions. Overall, we conclude that resetting of heritable silencing following stress is a regulated process which is mediated by the MAPK pathway and by the *skn-1* transcription factor.

To examine how stress-induced resetting affects the heritable small RNA molecules, we used small RNA sequencing. Endogenous small RNAs align to large parts of the genome, and their inheritance is especially important for transgenerational regulation of germline expressed transcripts (de Albuquerque et al., 2015; Gu et al., 2009; Phillips et al., 2015). We sequenced small RNAs from worms which were stressed at the L1 stage (starvation, heat shock, hyperosmotic conditions) and from their progeny (see scheme in **Figure 5A** and **Methods**). The sequencing results revealed that the overall levels of endogenous small RNAs decrease in response to each of the three stresses (See **Figure 5B** and **Methods**). Specifically, the levels of endogenous small RNAs which align in the antisense orientation to protein-coding genes, non-coding RNAs (ncRNAs), and pseudogenes were all significantly reduced following stress (**Figure 5B**).

**Figure 5:**
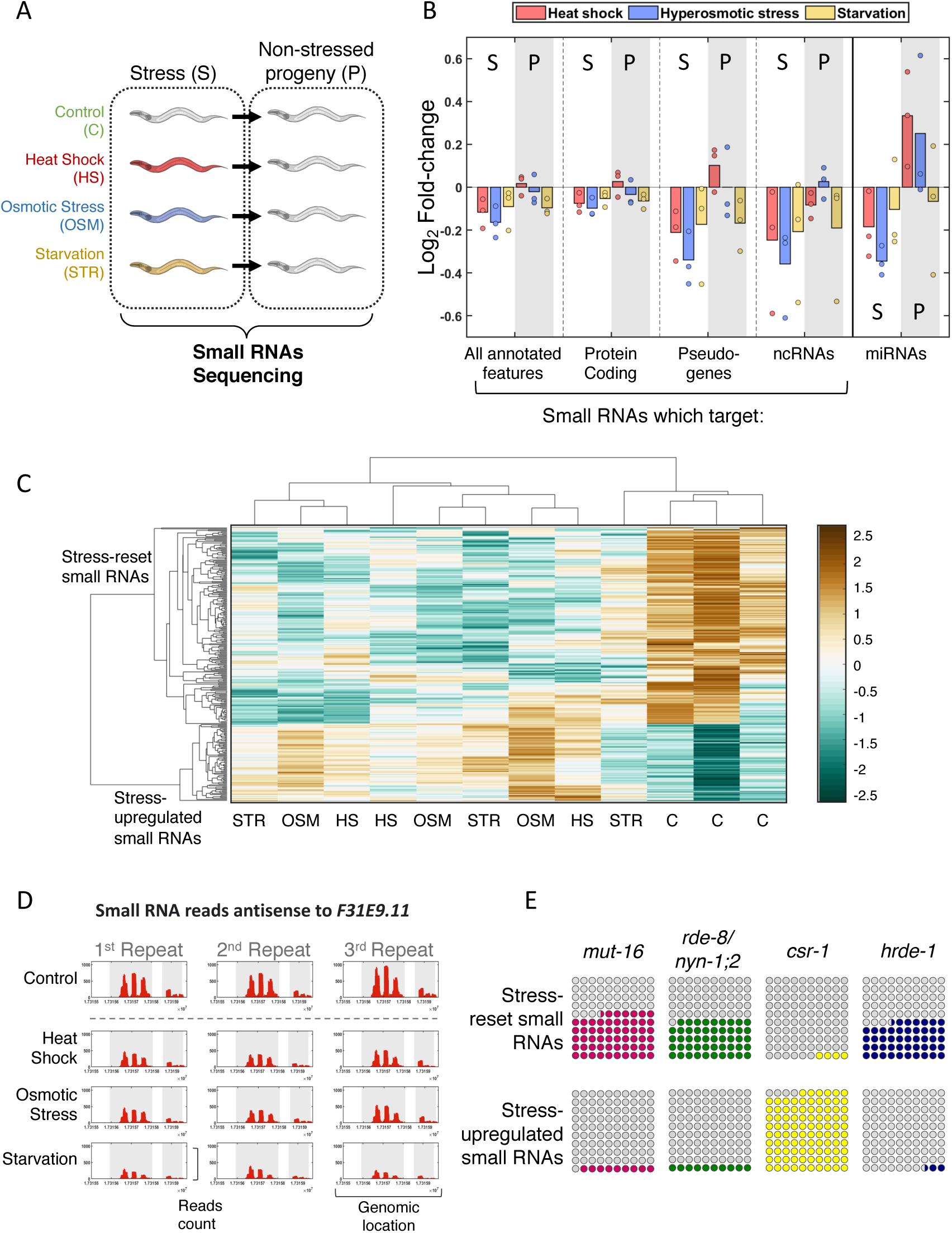
Genome-wide small RNA changes following stress. A. *Experimental scheme*. Worms were exposed to stress during their first larval stage and were collected for RNA extraction and small RNA sequencing on the first day of adulthood. The next generation was grown under normal conditions (See **Methods**). B. *Changes in total small RNA levels aligning to various genomic features.* Presented are the Log_2_ fold-change values (y-axis) for each condition (color coded) when compared to the control. Each dot represents one independent biological repeat. C. Clustering of the 281 stress-affected small RNA targets, based on their normalized total number of reads in each sample (4 different conditions in 3 independent biological repeats). Hierarchical clustering was performed with Spearman’s rank correlation as a distance metric and ‘average’ linkage. The data have been standardized across all columns for each gene, so that the mean is 0 and the standard deviation is 1. D. *An example of a stress-affected small RNAs target*. The *F31E9.11* gene is covered by small RNAs which are reset across all stress conditions. Shown are the normalized read counts (y-axis) as function of genomic location (x-axis) of small RNAs targeting the *F31E9.11* gene. Exons appear on a grey background. E. *Overlap of targets of stress-affected small RNAs with known targets of different small RNA pathways.* Each square represents the proportional overlap of reset (upper row) or up-regulated (bottom row) small RNA targets with known targets of the indicated small RNA pathways. Shown are results for MUT-16-dependent small RNA targets (Zhang et al., 2011), NYN-1;2/RDE-8-dependent small RNA targets (Tsai et al., 2015), CSR-1-bound small RNA targets (Claycomb et al., 2009) and HRDE-1-bound small RNA targets (Buckley et al., 2012).

In worms that were directly exposed to stress, all three types of stress conditions led to resetting of heritable endogenous small RNA levels. However, across multiple repeats (see **Methods**), different stressors shape the small RNA pools of the next generation in different ways (**Figure 5B**). Non-stressed progeny that derive from starved parents maintained the reduction in the overall levels of the different types of endogenous small RNA in all three repeats. In contrast, in non-stressed progeny that were derived from heat-shocked parents the levels of most types of endogenous small RNAs were elevated. We did not detect a consistent heritable change in the global levels of endogenous small RNAs in non-stressed progeny that derive from parents that experienced hyperosmotic stress (**Figure 5B**).

When comparing all samples derived from stressed worms, we identified a list of 281 genes that were targeted by stress-affected endogenous small RNAs with high confidence (FDR<0.1, **Figure 5C, 5D** and **Supplementary Table 1**). In the non-stressed progeny of these worms, we did not detect wide-spread changes in stress-affected small RNAs (only 10 genes were targeted by such small RNAs, **Supplementary Table 2**). In 72% (202/281) of the gene targets of stress-affected small RNAs, the targeting small RNAs were down-regulated, or “reset”, following stress. The small RNAs which target the remaining 79 (28%) genes were up-regulated following stress and were almost exclusively CSR-1-dependent endo-siRNAs (hypergeometric test, p-value=2.9 e^−48^, 96% were found to be physically bound to CSR-1, (Claycomb et al., 2009)) (**Figure 5E**). Unlike other endo-siRNAs, CSR-1 endo-siRNAs were demonstrated to promote gene expression rather than gene silencing (Wedeles et al., 2013). The endogenous small RNAs that were down-regulated following stress exhibited a strong enrichment for mutator-dependent small RNAs (FDR= 2.2e^−75^, (Phillips et al., 2014; Yang et al., 2016; Zhang et al., 2011)); Small RNAs which are reset following stress overlap with small RNAs that are depleted due to defects in proteins which are localized to the mutator foci, granules that are required for small RNA amplification (Phillips et al., 2012) (see **Figure 5C** and **5E**). These proteins (*mut-16*, *mut-14*;*smut-1* (Phillips et al., 2014; Zhang et al., 2011)) were shown to be involved in multiple endogenous small RNA biogenesis pathways, affecting both somatic and germline small RNAs, and are required for efficient small RNA amplification and cleavage of target RNAs (*rde-8* and *nyn-1;nyn-2* (Tsai et al., 2015)). Interestingly, generation of *mut-14;smut-1*;*mut-16* triple mutants was shown to enable erasure of heritable small RNA-based memory of self and non-self genes (de Albuquerque et al., 2015; Phillips et al., 2015). We further found that targets of stress-reduced small RNAs are enriched for dsRNA-producing loci (FDR=0.0019) (Saldi et al., 2014) and for piRNAs gene targets (FDR= 3.2e^−07^) (Bagijn et al., 2012) (See full list of enrichments in **Supplementary Table 3**). Out of the gene targets of stress-affected small RNAs, 31/281 (FDR=2.8e^−14^) were previously identified as genes which are targeted by small RNAs transgenerationally in response to L1 starvation (Rechavi et al., 2014). Further, the 281 genes targeted by stress-affected small RNAs show enrichment for stress-related functions. As a group, these genes were shown to be regulated by the TGF-beta pathway (FDR=2.4e^−08^) (Shaw et al., 2007), the insulin pathway (FDR=1.2e^−08^) (Halaschek-Wiener et al., 2005), and to be affected in response to rapamycin exposure (FDR=0.0023 (Calvert et al., 2016)), and changes in temperature (FDR=0.0027 (Viñuela et al., 2011)). Specifically, we found a significant enrichment for target genes which are regulated by SKN-1 and by KGB-1 (FDR=0.00095 and FDR= 0.032 respectively), which we identified as regulators of stress-induced resetting of small RNAs (**Figure 4C and Supplementary Figure 2**). Overall, our sequencing data show that different types of stress dramatically reshape the heritable small RNA pool, affecting in particular stress-related targets.

In a previous study, we found that MET-2, a methyltransferase required for mono- and di-methylation of H3K9, is essential for termination of RNAi inheritance, and that in *met-2* mutants aberrant endo-siRNAs accumulate over generations, leading eventually to sterility (Mortal Germline, Mrt phenotype) (Lev et al., 2017). Interestingly, the targets of stress-affected small RNAs were enriched for genes which were previously found to be mis-regulated in *met-2* mutants (FDR=2.1e-05) (Zeller et al., 2016) and we found that these genes have exceptionally high levels of H3K9me2 (**Figure 6A, Supplementary Figure 4A** and **Methods**).

**Figure 6:**
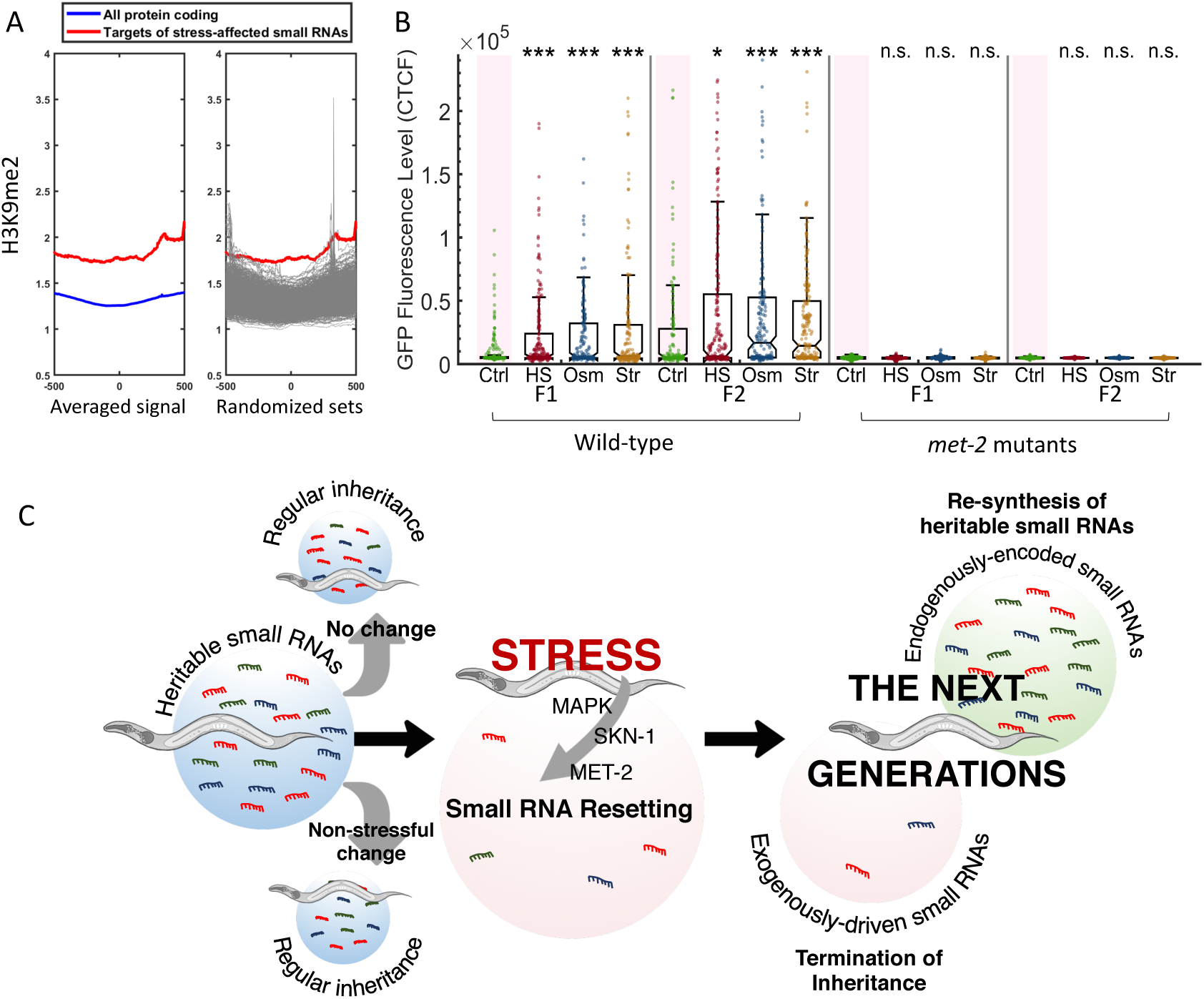
Stress-induced small RNA resetting depends on the H3K9 methyltransferase MET-2. A. *Targets of stress-affected small RNAs show significantly increased H3K9me2 marks*. An analysis of *H3K9me2* signal (based on published data from (McMurchy et al., 2017)). Presented is the averaged H3K9me2 signal (y-axis) of all protein coding genes (blue) and target genes of stress-affected small RNAs (red). All genes are aligned according to their Transcription Start Sites (TSS), and the regions of 500 base pairs upstream and downstream of the TSS are shown on the x-axis. Each gray line (right panel) represents the average of a random set of genes (500 iterations) equal in size to the set of target genes of stress-affected small RNAs. B. *met-2 mutant worms do not reset heritable small RNAs in response to stress*. The graph displays the measured germline GFP fluorescence levels of wild-type and *met-* 2 mutant worms (y-axis) across generations under the indicated condition (x-axis). Each dot represents the value of an individual worm. Shown are the median of each group, with box limits representing the 25^th^ (Q1) and 75^th^ (Q3) percentiles, notch representing a 95% confidence interval, and whiskers indicating Q1-1.5*IQR and Q3+1.5*IQR. FDR-corrected values were obtained using Dunn’s test. Not significant (ns) indicates q ≥ 0.05, (*) indicates q<0.05, and (***) indicates q<0.001, see **Methods**). C. *A model summarizing stress-induced resetting of heritable small RNAs.* Small RNAs from both endogenous and exogenous sources are reset in response to stress. Resetting is mediated by the MAPK pathway, the SKN-1 transcription factor and the H3K9 methyltransferase MET-2. Endogenous small RNAs which are encoded in the genome are re-synthesized in the next generations, while small RNAs from exogenous sources and transient responses are eliminated.

We therefore examined if MET-2’s regulation over RNAi inheritance is required also for the resetting that follows exposure to stress. We found that MET-2 is essential for stress-induced resetting of heritable RNAi (**Figure 6B**). Further, even when we examined inheritance of weak RNAi responses (by tittering down the amount of dsRNA used to induce RNAi), *met-2* mutants were resistant to small RNAs resetting (**Supplementary Figure 4B**). Thus, MET-2 is required for the execution of stress-induced small RNA resetting, even when accounting for the exceptionally strong RNAi inheritance in *met-2* mutants.

In summary, we found that worms erase heritable small RNAs in response to stress, and that stress response pathways affect the dynamics of RNAi inheritance. Resetting is an actively regulated process which depend on the MET-2 methyltransferase. This mechanism can be important for ensuring heritable small RNAs maintain environment-relevant genes regulation programs.

## Discussion

In this study we found that stress leads to resetting of transgenerationally transmitted small RNAs, erasing heritable gene regulatory responses. Stress-induced resetting depends on the MAPK pathway and the SKN-1 transcription factor, and is regulated by MET-2, an H3K9 methyltransferase that is required for germline reprogramming (Kerr et al., 2014). We found that stress induces a reduction in small RNA levels across the genome. This reduction stems from changes in multiple distinct small RNA species, including small RNAs that regulate stress-related genes. We speculate that a mechanism for resetting of heritable small RNAs could be adaptive in rapidly changing environments (See model in **Figure 6C**).

We previously described a tunable mechanism that controls the duration of heritable RNAi responses and found that ancestral RNAi responses can be enhanced by non-target-specific reactivation of the RNAi system in the progeny (Houri-Ze’evi et al., 2016). We hypothesized that when both parents and progeny are challenged by dsRNA-induced RNAi it could be beneficial for the worms to extend the duration of ancestral RNAi responses. We reasoned this could be the case since under these circumstances the child’s environment resembles the parent’s, and therefore the heritable response could still be relevant for the progeny. In contrast, stress-induced resetting of heritable small RNAs, the phenomenon described in this manuscript, could unburden descendants from heritable information that is no longer relevant.

Accumulating evidence from the past two decades, collected in different organisms, suggest that stress can lead to intergenerational and transgenerational epigenetic changes (Bohacek and Mansuy, 2015). In certain instances, stress was found to lead to transient heritable reduction in small RNAs and chromatin marks (Belicard et al., 2018; Klosin and Lehner, 2016). Such heritable effects on chromatin were referred to as “epigenetic wounds” which “heal” following gradual re-accumulation of the marks over generations (Klosin and Lehner, 2016). Our results suggest that reduction in small RNAs following stress is an active and regulated process which is initiated by stress-signaling pathways (and specifically, the MAPK pathway).

Active removal of epigenetic regulation in response to stress could serve as a mechanism for increasing genetic and phenotypic variability. Small RNAs in *C.elegans* distinguish between self and non-self genes (Shirayama et al., 2012), and orchestrate gene expression in the germline and during development (Feng and Guang, 2013; Gu et al., 2009; Han et al., 2009). Relieving the regulation of heritable small RNAs could be a way to increase phenotypic plasticity or to reveal hidden genetic variability. For example, a conserved microRNA in flies was shown to buffer developmental programs against variation (Li et al., 2009).

We have recently found that changes in neuronal small RNA levels generate transgenerational effects (Posner et al., in Press). Similarly, it has been shown that olfactory memory can become heritable (Moore et al., 2018; Pereira et al., 2019; Remy, 2010). The systemic response to stress that KGB-1, SKN-1 and SEK-1 mediate (factors which are shown here to be required for resetting) was shown in the past to be coordinated by the nervous system (Bishop and Guarente, 2007; Liu et al., 2018; Shivers et al., 2009). Therefore, it could be interesting to understand if and how resetting of heritable small RNAs is controlled by neuronal activity.

Non-DNA-based inheritance could be involved in multiple human disorders (Crews et al., 2014; Nilsson et al., 2018; Tucci et al., 2019). It is still unclear whether the same mechanisms that allow transgenerational inheritance in worms exist in mammals (Horsthemke, 2018). If analogous mechanisms are indeed conserved, understanding the pathways that counteract transmission or maintenance of heritable small RNAs could aid in prevention or treatment of various diseases.

## Acknowledgments

We thank all members of the Rechavi lab for fruitful discussions and for their support. Some strains were provided by the CGC, which is funded by NIH Office of Research Infrastructure Programs (P40 OD010440). We thank the Richard Roy lab for providing strain MR1175 (*aak-1/2*) and the Julie Ahringer lab for providing strain JA1527. We thank Yoav Ze’evi (statistics unit, Yoav Benjamini’s group) for his help with the statistical analysis. O.R is thankful to the Adelis foundation grant #0604916191. L.H-Z. is thankful to the Clore foundation. The Rechavi lab is funded by ERC grant #335624 and the Israel Science Foundation (grant#1339/17).

## Supplementary Figures

**Supplementary Figure 1:**
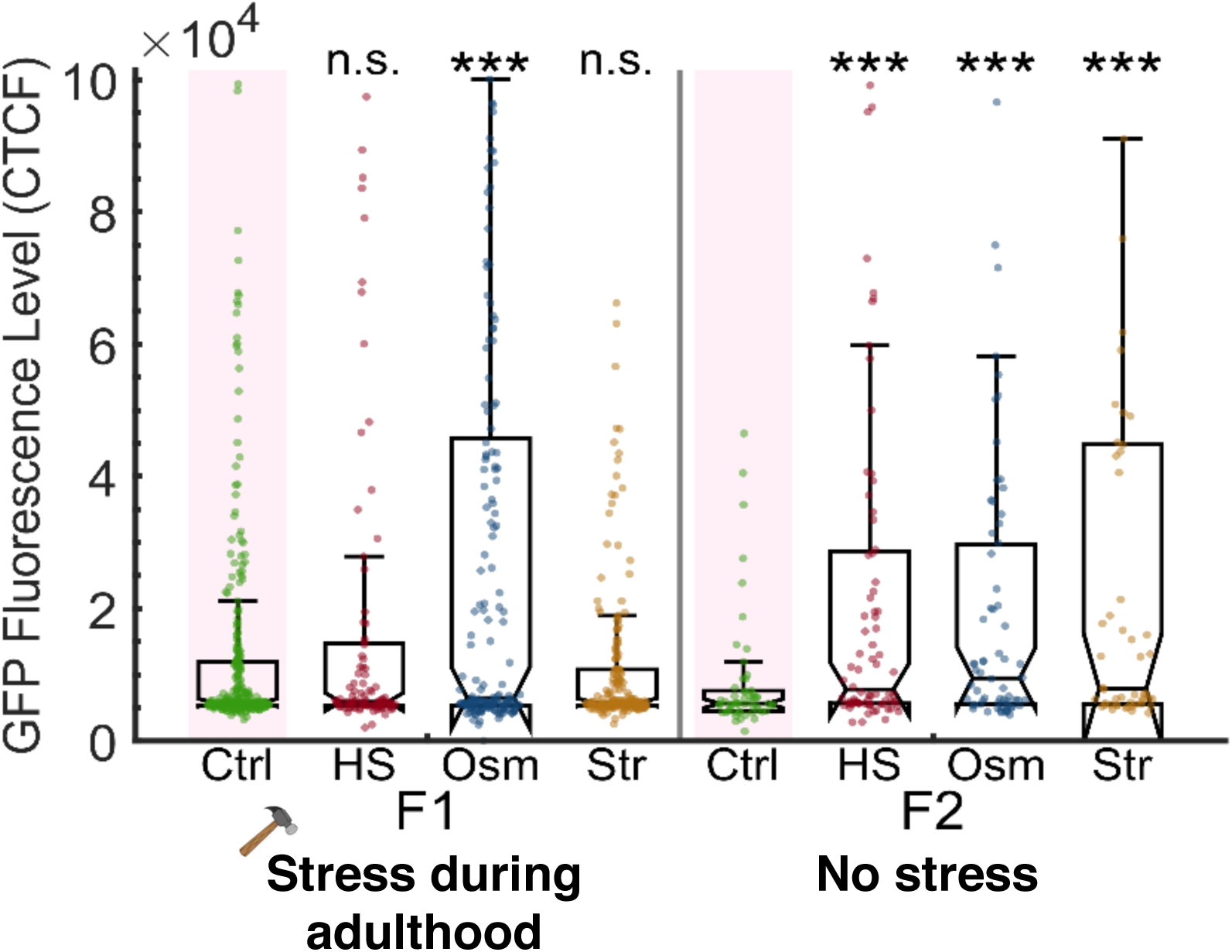
Stress resets heritable small RNAs even when applied during adulthood. Related to Figure 1. *Stress applied during adulthood leads to resetting of heritable silencing in the next generation.* The graph displays the measured germline GFP fluorescence levels of individual worms (y-axis) across generations under the indicated condition (x-axis). Each dot represents the value of an individual worm. Shown are the median of each group, with box limits representing the 25^th^ (Q1) and 75^th^ (Q3) percentiles, notch representing a 95% confidence interval, and whiskers indicating Q1-1.5*IQR and Q3+1.5*IQR. FDR-corrected values were obtained using Dunn’s test. Not significant (ns) indicates q ≥ 0.05 and (***) indicates q<0.001 (see **Methods**).

**Supplementary Figure 2:**
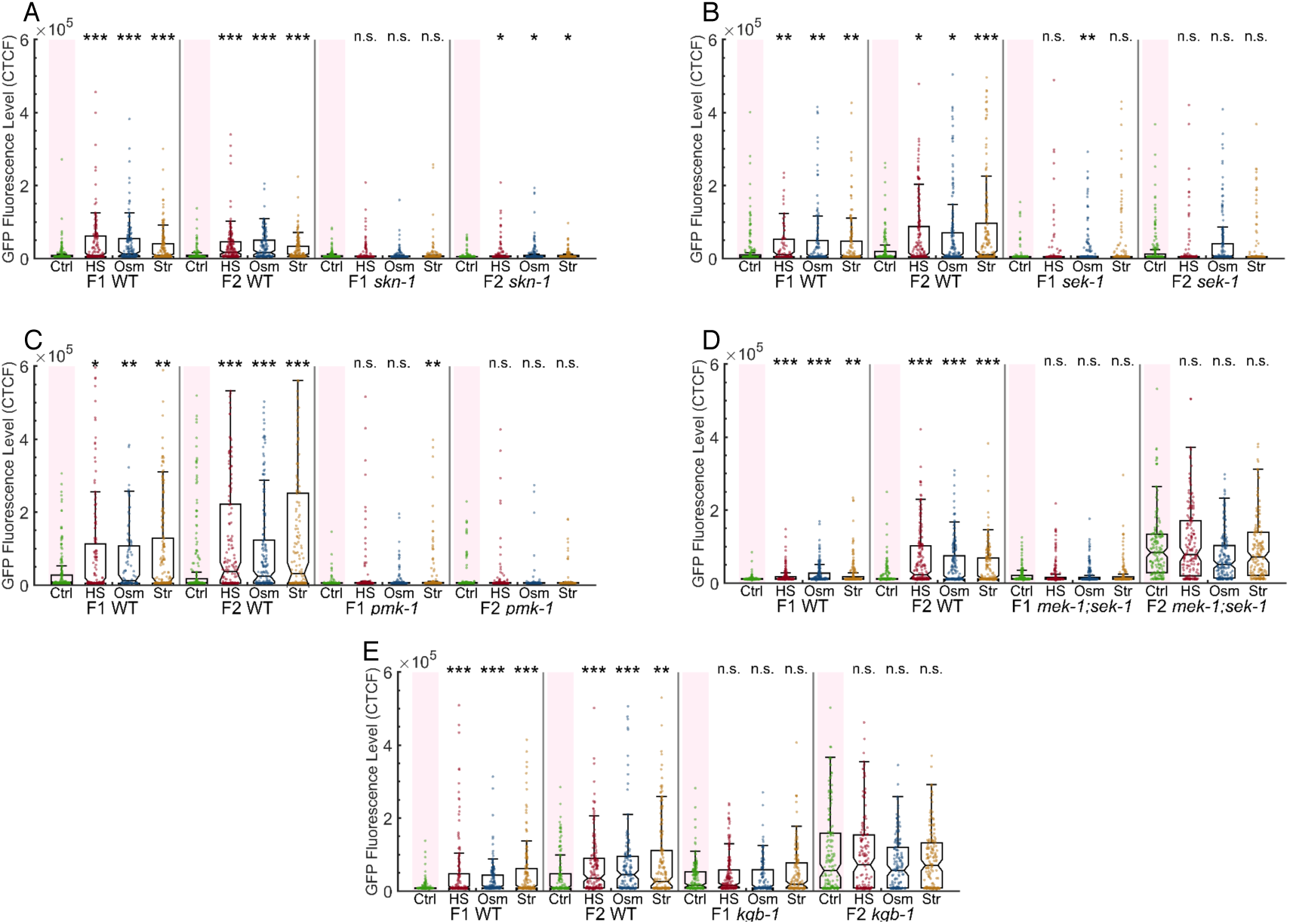
Multiple stress signaling and processing pathways affect heritable RNAi dynamics and stress-induced resetting of heritable silencing. Related to Figure 4. The graphs display the measured germline GFP fluorescence levels of mutant worms (y-axis) across generations under the indicated condition (x-axis). Each dot represents the value of an individual worm. Each mutant was examined in an independent experiment. Shown are the median of each group, with box limits representing the 25^th^ (Q1) and 75^th^ (Q3) percentiles, notch representing a 95% confidence interval, and whiskers indicating Q1-1.5*IQR and Q3+1.5*IQR. FDR-corrected values were obtained using Dunn’s test. Not significant (ns) indicates q ≥ 0.05, (*) indicates q<0.05, (**) indicates q<0.01, and (***) indicates q<0.001, see **Methods**). Each mutant strain was examined side-by-side with WT worms.

i. *sek-1* mutants do not reset a heritable silencing response following stress
ii. *pmk-1* mutants do not reset a heritable silencing response following stress
iii. *mek/sek-1* mutants do not reset a heritable silencing response following stress
iv. *kgb-1* mutants do not reset a heritable silencing response following stress
v. *skn-1* hypomorphs do not reset a heritable silencing response following stress

**Supplementary Figure 3:**
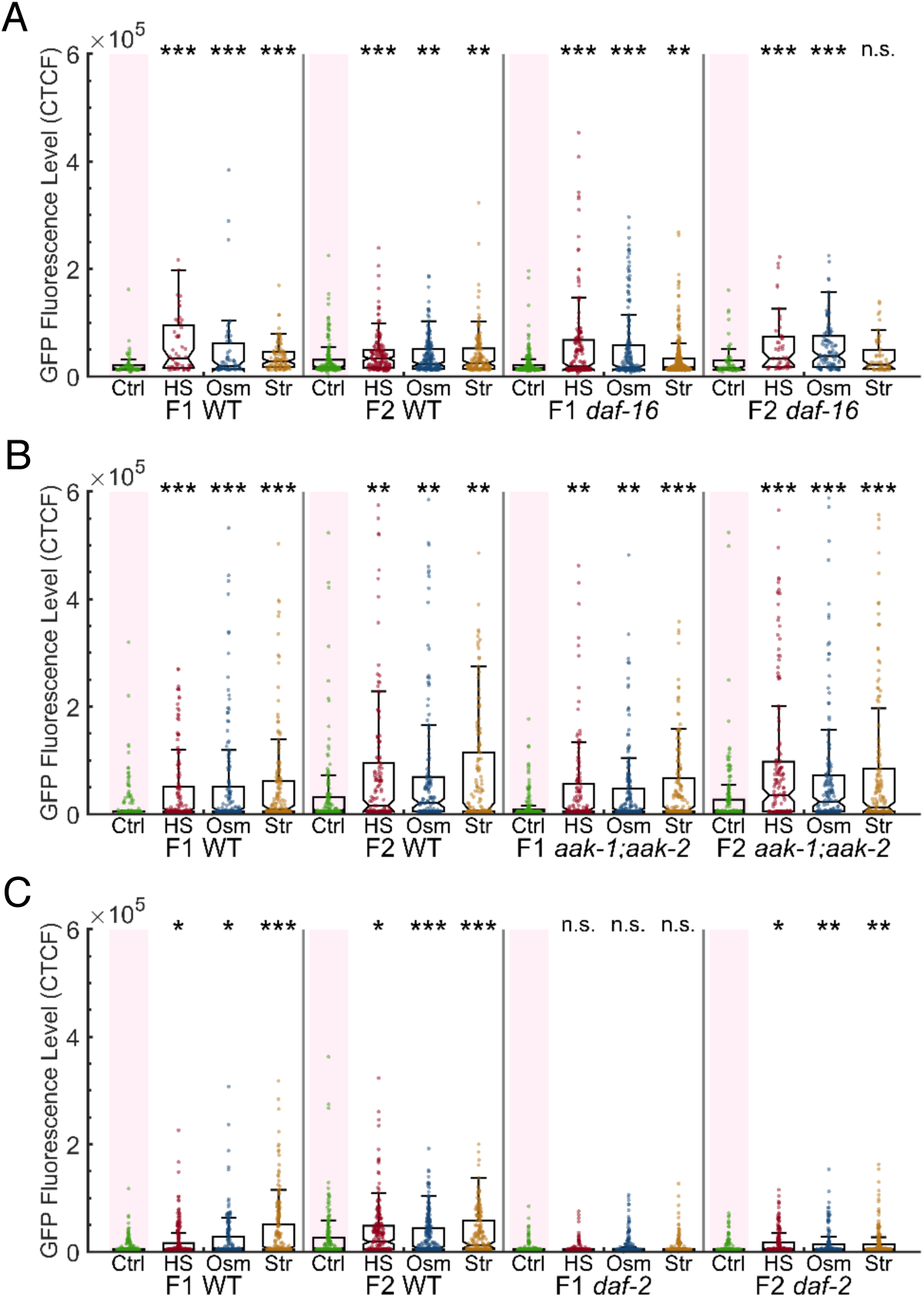
Mutations in daf-16, aak-1/2 and daf-2 genes do not affect stress-induced resetting of heritable responses. Related to Figure 4. The graphs display the measured germline GFP fluorescence levels of mutant worms (y-axis) across generations under the indicated condition (x-axis). Each dot represents the value of an individual worm. Each mutant was examined in an independent experiment. Shown are the median of each group, with box limits representing the 25^th^ (Q1) and 75^th^ (Q3) percentiles, notch representing a 95% confidence interval, and whiskers indicating Q1-1.5 *IQR and Q3+1.5*IQR. FDR-corrected values were obtained using Dunn’s test. Not significant (ns) indicates q ≥ 0.05, (*) indicates q<0.05, (**) indicates q<0.01, and (***) indicates q<0.001, see **Methods**). Each mutant strain was examined side-by-side with WT worms.

i. *daf-16* mutants show regular RNAi inheritance dynamics and reset a heritable silencing response following stress.
ii. *aak-1/2* mutants show regular RNAi inheritance dynamics and reset a heritable silencing response following stress.
iii. *daf-2* mutants show *enhanced* RNAi inheritance dynamics and reset a heritable silencing response following stress.

**Supplementary Figure 4:**
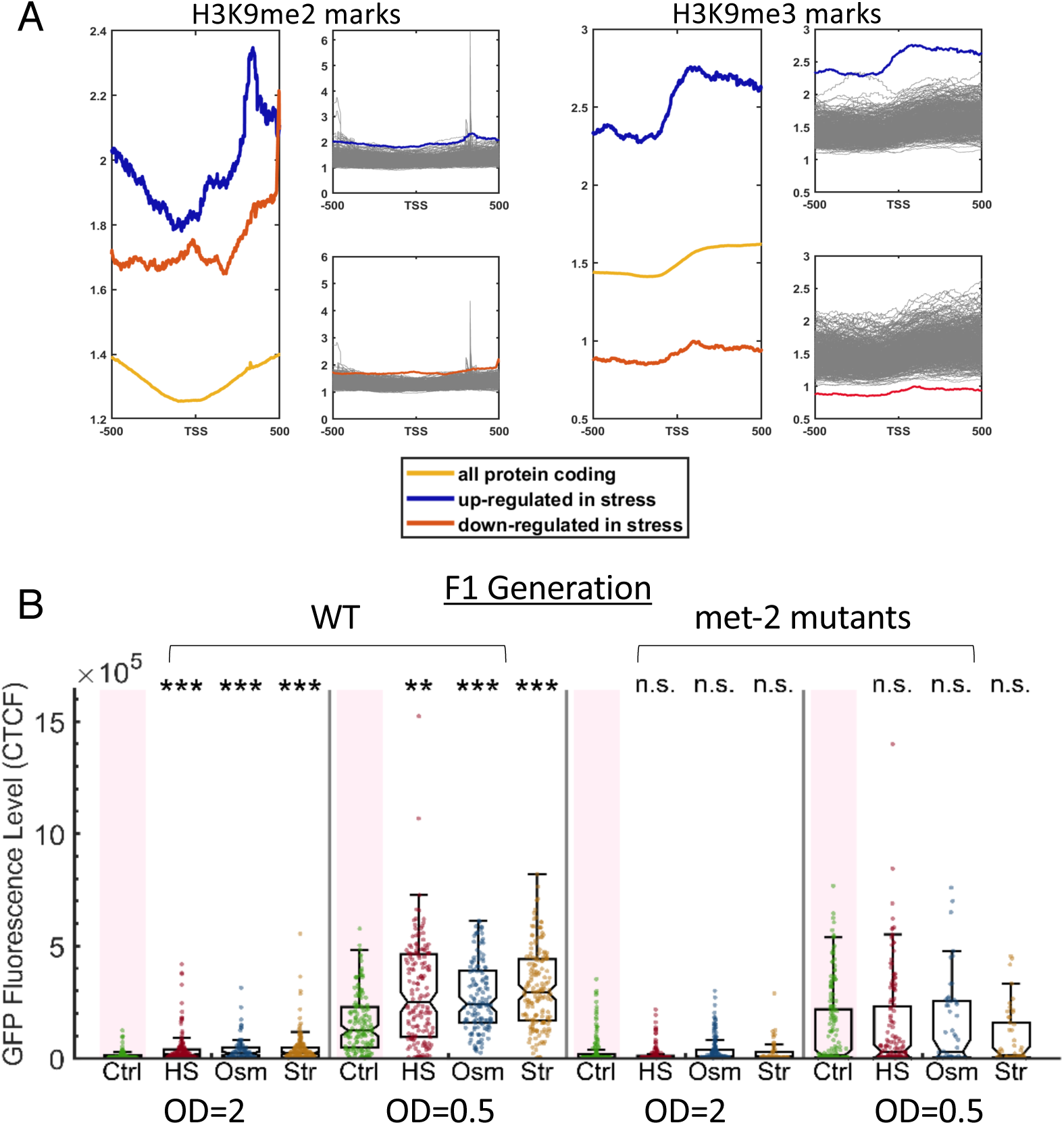
Targets of stress-affected small RNAs show unique H3K9 marks and the H3K9 methyltransferase met-2 is required for the execution of small RNA resetting. Related to Figure 6. A. *Targets of stress-affected small RNAs show significantly different H3K9me2 and H3K9me3 marks*. An analysis of *H3K9me2* and H3K9me3 signals (based on published data from (McMurchy et al., 2017)). All genes are aligned according to their Transcription Start Sites (TSS), and the regions of 500 base pairs upstream and downstream of the TSS are shown on the x axis. The y axis shows the averaged signal of the H3K9me2 (left panel) and H3K9me3 (right panel) modifications as a function of distance from the TSS. The chromatin modification profile is shown for 3 genes sets: all protein coding (yellow), set of genes that are up-regulated in stress (blue) and set of genes that are down-regulated in stress (red). Each gray line represents the average of a random set of genes equal in size to the set of up-regulated (upper panels) or down-regulated (lower panels) genes. B. *met-2 mutants do not reset heritable silencing even when weaker RNAi inheritance responses are induced*. WT and *met-2* worms are exposed at the P0 generation to dsRNA-producing bacteria of varying concentrations (either OD=2 or OD=0.5) and examined for stress-induced resetting at the F1 generation. The graphs display the measured germline GFP fluorescence levels of WT and *met-2* mutant worms (y-axis) under the indicated condition (x-axis). Each dot represents the value of an individual worm. Shown are the median of each group, with box limits representing the 25^th^ (Q1) and 75^th^ (Q3) percentiles, notch representing a 95% confidence interval, and whiskers indicating Q1-1.5*IQR and Q3+1.5*IQR. FDR-corrected values were obtained using Dunn’s test. Not significant (ns) indicates q ≥ 0.05, (**) indicates q<0.01, and (***) indicates q<0.001, see **Methods**).

## Materials and Methods

### EXPERIMENTAL PROCEDURES

#### Worms’ maintenance

Standard culture techniques were used to maintain the nematodes. The worms were grown on Nematode Growth Medium (NGM) plates and fed with OP50 bacteria. Except when otherwise noted, all experiments were performed at 20°C. The worms were kept fed for at least five generations before the beginning of each experiment. Extreme care was taken to avoid contamination or starvation. Contaminated plates were discarded from the analysis. All experiments were performed with at least 3 biological replicates. The nematode strains used in this study are indicated in the Key Resources Table.

#### RNAi treatment

The standard assay for RNAi by feeding was carried out as previously described (Kamath et al., 2000): HT115 bacteria that transcribe dsRNA targeting *gfp*, or control HT115 bacteria that only contain an empty vector plasmid, were grown in LB containing Carbenicillin 1:1000, and were than seeded on NGM plates that contain Carbenicillin 1:1000 and IPTG 1:1000. In the generation of worms that was exposed to RNAi (P0), worms were cultivated on those plates.

In experiments where the concentration of RNAi bacteria was controlled, a spectrophotometer was used to measure optical density at OD600. HT115 bacteria were diluted with empty vector bacteria to 2 OD and 0.5 OD.

#### L1 stress

Synchronized eggs were obtained by bleaching gravid adult worms. The progeny was then subjected to the various stress conditions.

##### Heat shock

Heat shock treatment was performed at 37°C. The nematodes were heat shocked 24 hours after the bleach treatment, for a duration of 120 minutes. After heat shock, the treated worms were placed back in an incubator set to 20°C, with each plate placed in a separate location (not in a pile) to avoid temperature differences between top/bottom plates and the middle plates.

##### Hyperosmotic stress

After bleach, the obtained eggs were seeded on high-salt growth media plates (350mM NaCl) and were grown on these plates for 48 hours. After 48 hours, the worms were washed off with M9 buffer onto regular NGM plates.

##### Starvation

For starvation conditions, the nematodes were grown on empty NGM plates and were maintained this way for a total of 6 days. After 6 days, the worms were washed off with M9 onto regular NGM plates seeded with OP50 bacteria. In the experiment with *aak-1/2* double mutants, both the wild-type and mutant worms were starved for 3 days instead of 6 days, due to starvation-sensitivity of the mutants.

In the experiments described in **Figure 2b** (a reverse change in settings; from stressed conditions to regular growth conditions), P0 nematodes were grown to adulthood on plates with RNAi bacteria in either a 20°C or a 25°C. The nematodes were bleached in adulthood, and their progeny were grown to adulthood on NGM plates either in a 20°C or a 25°C.

#### Adults stress

In the adult stress assays (**Supplementary Figure 1b**), stress conditions were applied when the worms reached young adulthood, with heat shock lasting one hour, hyperosmotic stress lasting 24 hours, and starvation lasting 48 hours.

#### The next generations

In both assays (L1 and adults stress), the next generation of worms was cultured by randomly picking several mothers from each plate to a fresh plate and allowing them to lay eggs for few hours (8-24 hours). The number of picked worms and length of egg-laying period were constant within each experiment across all plates and conditions.

#### piRNAs- and endo-siRNAs-derived silencing assays

To obtain expression of the piRNAs-silenced mCherry and the endo-siRNAs-silenced GFP, worms were grown in 25°C for 3 generations and then transferred back to recovery at 20°C. After two generations, worms were bleached and assayed for the different stress conditions, as described above.

#### Fluorescence microscopy

We used an *Olympus BX63* microscope for fluorescence microscopy assays. Experiments were filmed with a 10X objective lens, with an exposure time of 750ms.

##### Small RNAs sequencing

###### Collecting the worms

Total RNA was extracted from “day one” adults. As stress induces variability in development even in isogenic worms’ populations, tight synchronization of the worms’ populations was achieved by picking L4 worms one day prior to collecting the worms for sequencing. Each experiment started by exposure of the generation before stress to an anti-*gfp* RNAi trigger to enable the phenotypic detection of resetting in the next generations. All sequencing experiments were done in triplicates (independent biological replicates).

###### Libraries preparation

Worms were lysed using the TRIzol® reagent (Life Technologies) followed by repetitive freezing, thawing, and vortex. The total RNA samples were treated with tobacco acid pyrophosphatase (TAP, Epicenter), to ensure 5′ monophosphate-independent capturing of small RNAs. Libraries were prepared using the NEBNext® Small RNA Library Prep Set for Illumina® according to the manufacturer’s protocol. The resulting cDNAs were separated on a 4% agarose E-Gel (Invitrogen, Life Technologies), and the 140–160 nt length species were selected. cDNA was purified using the MinElute Gel Extraction kit (QIAGEN). Libraries were sequenced using an Illumina NextSeq500 instrument.

### QUANTIFICATION AND STATISTICAL ANALYSIS

#### Fluorescence analysis; GFP

Using the ImageJ *Fiji* “measure” function (Schindelin et al., 2012), we measured the integrated density of the three germline nuclei closest to spermatheca in each worm, as well as a mean background measurement of the worm in the germline’s vicinity. If less than three germline nuclei were visible, a measurement was taken in the estimated location of the germline instead. The corrected total cell fluorescence (CTCF) of each germline nucleus was calculated <u>as previously described</u>. We used the mean of all three CTCF scores as the worm’s fluorescence score.

#### Fluorescence analysis; piRNAs- and endo-siRNAs-derived silencing

Using *Fiji* (Schindelin et al., 2012), we have measured the integrated density of the whole worm, as well as one background measurement per worm. The corrected total fluorescence (CTF) of each worm was calculated as described above.

#### Statistical analyses

Due to the highly variable nature of small RNA inheritance and erasure, we have elected to analyze our data using two - tailed Dunn’s multiple comparison test. We corrected for multiple comparisons using the Benjamini-Hochberg false discovery rate (FDR), with an FDR of 0.05. The q-values reported in this study are adjusted to multiple comparisons. In the figures, not significant (ns) indicates q ≥ 0.05, (*) indicates q<0.05, (**) indicate q<0.01, and (***) indicate q<0.001. Unless marked otherwise in the figures, comparisons were made between each stressed group and its corresponding unstressed control group.

#### Box plot graphs

Data are the individual worms’ mean germline CTCF, with each dot representing the measurement of a single worm. Data are represented as medians, with box limits representing the 25^th^ (Q1) and 75^th^ (Q3) percentiles, notch representing a 95% confidence interval, and whiskers indicating Q1-1.5*IQR and Q3+1.5IQR. The minimal value of each experiment was set at 0 and the other values were adjusted accordingly.

#### Small RNA-seq Analysis

The Illumina fastq output files were first assessed for quality, using FastQC (Andrews, 2010), and compared to the FastQC-provided example of small RNA sequencing results. The files were then assigned to adapters clipping using Cutadapt (Martin, 2011) and the following specifications were used:

cutadapt -m 15 -a AGATCGGAAGAGCACACGTCT input.fastq > output.fastq

**-m 15**: discard reads which are shorter than 15 nucleotides after the adapter clipping process

**-a AGATCGGAAGAGCACACGTCT**: the 3’ adapter sequence used as a query The clipped reads were then aligned against the *ce11* version of the *C. elegans* genome using ShortStack (Shahid and Axtell, 2014) using the default settings:

ShortStack --readfile Input.fastq –genomefile Ce11Reference.fa Next, we counted reads which align in the sense or antisense orientation to genes. Since stress is known to affect the abundance of structural RNA molecules, we omitted reads which align to structural genes from our analyses. We used the python-based script HTSeq-count (Anders et al., 2014) and the Ensembl-provided gff file (release-95), using the following command:

##### Antisense

HTSeq.scripts.count --stranded=reverse --mode=union input.sam GENES.gff > output.txt

##### Sense

HTSeq.scripts.count --stranded=yes --mode=union input.sam GENES.gff > output.txt

We then assigned the summarized counts for differential expression analysis using the R package DESeq2 (Love et al., 2014) and limited the hits for genes which were shown to have an FDR<0.1. Normalization of the total number of reads in each sample, and the total number of reads which align to the different types of genomic features (**Figure 5b**) was generated based on the SizeFactor normalization provided by the DESeq2 package (the median ratio method).

#### Plotting H3K9me2 and H3K9me3 profiles

Chip-Seq data were downloaded from GSE87522. The transcription start site (TSS) of all the protein coding genes in *C. elegans* were extracted from UCSC Genome Browser. We aligned all the genes according to their Transcription Start Sites (TSS) and extracted the signal along the flanking regions of 500 base pairs upstream and downstream of the TSS. In the case of genes with two transcripts or more, we averaged the histone modification signal of all the corresponding transcripts. The signal of individual genes was determined as the averaged signal of the two published replicates of each chromatin modification.

## DATA AVAILABILITY

Description: Small RNAs sequencing Fastq and count files.

Sequencing and count files have been deposited in the GEO under ID code GSE129988

### Supplementary Tables

1. **Supplementary Table 1**: 281 targets of small RNAs which are affected across all stress conditions *at the stress generation*. Table presents DESeq2 comparison of Control vs. Stress samples. Related to **Figure 5**.
2. **Supplementary Table 2**: 10 targets of small RNAs which are affected across all stress conditions *at the next generation*. Table presents DESeq2 comparison of Control vs. Stress samples. Related to **Figure 5**.
3. **Supplementary Table 3**: Enrichment table for the 281 targets of stress affected genes, generated using WormExp (Yang et al., 2016). Related to **Figure 5**.

## KEY RESOURCES TABLE

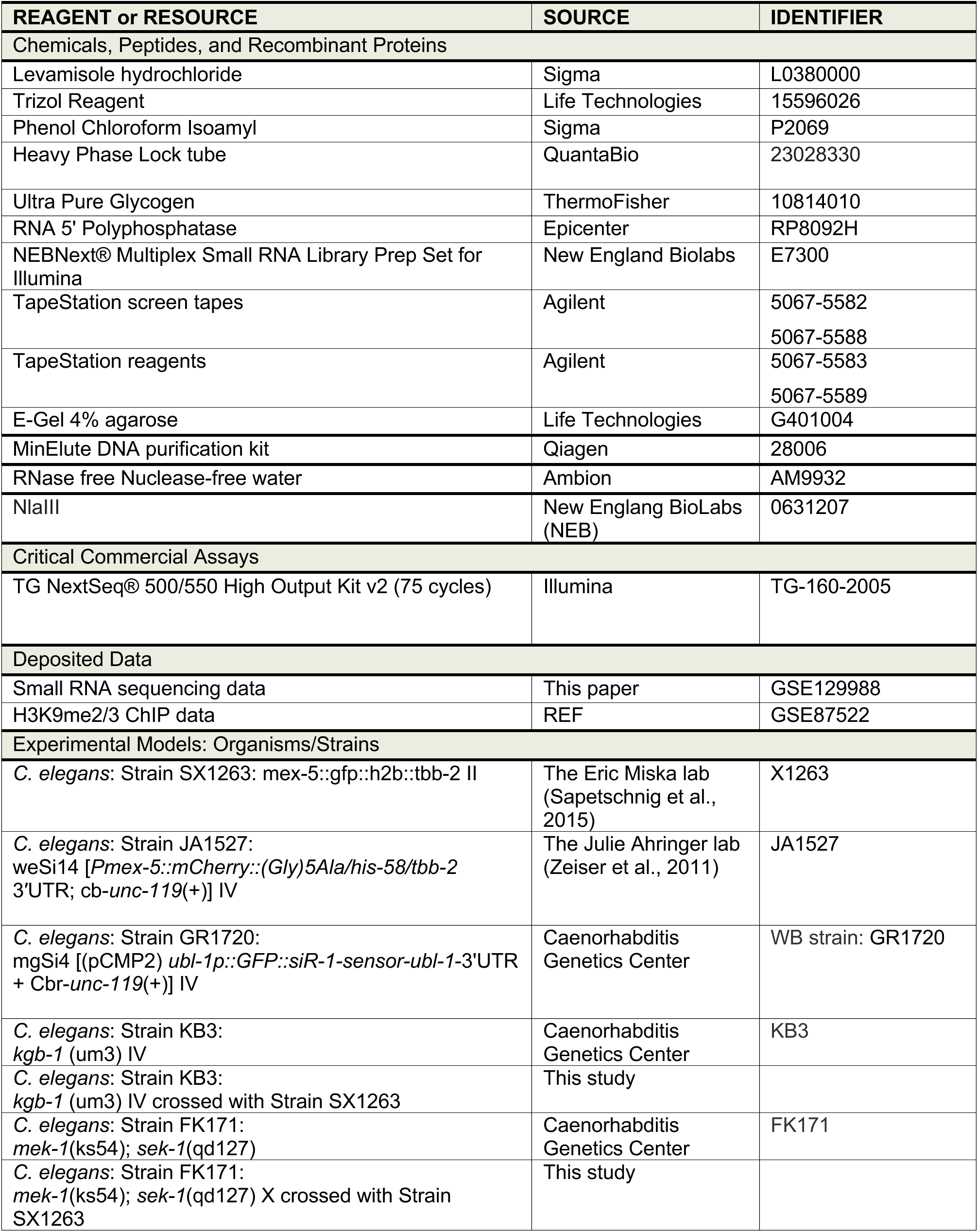
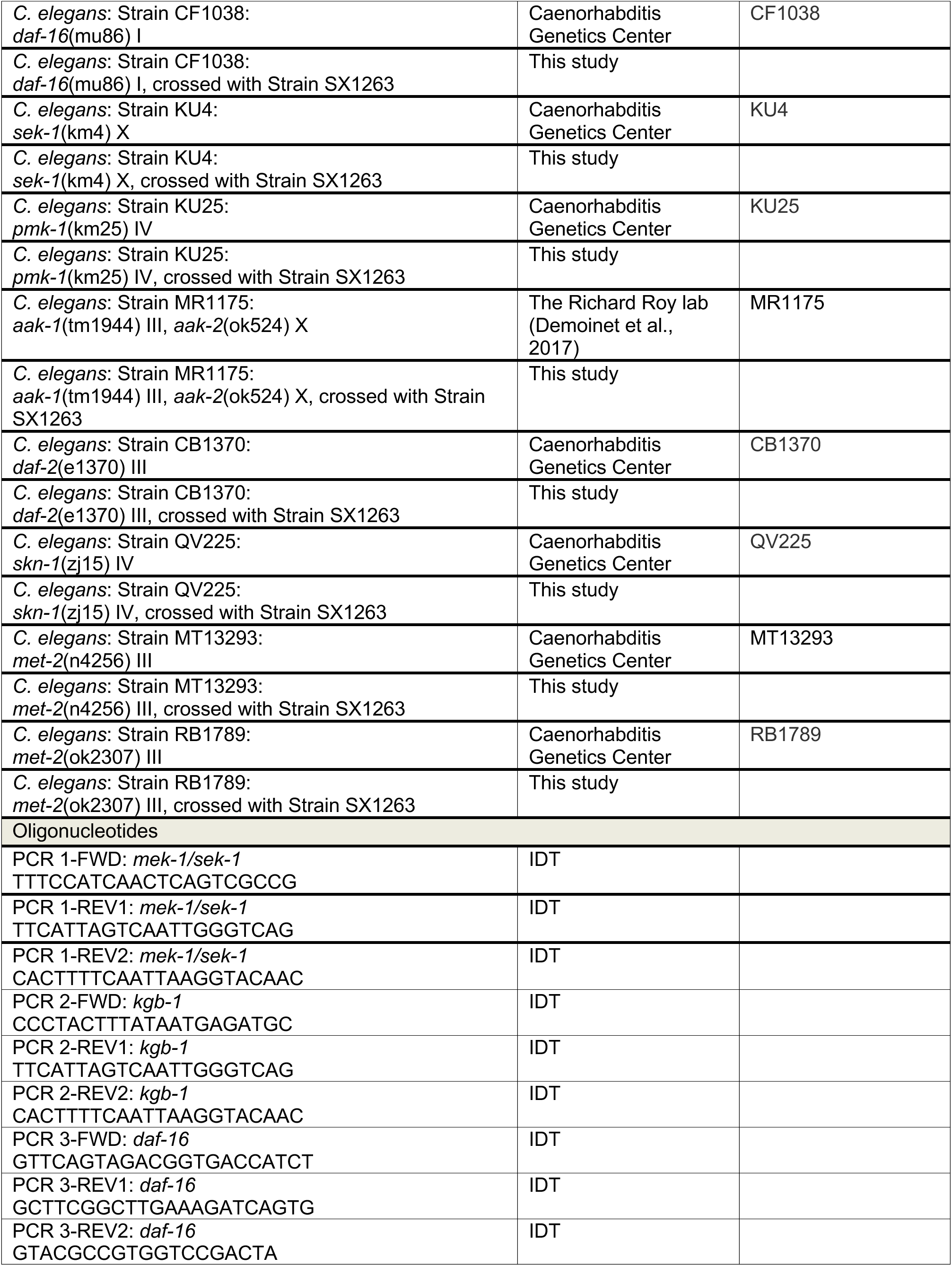
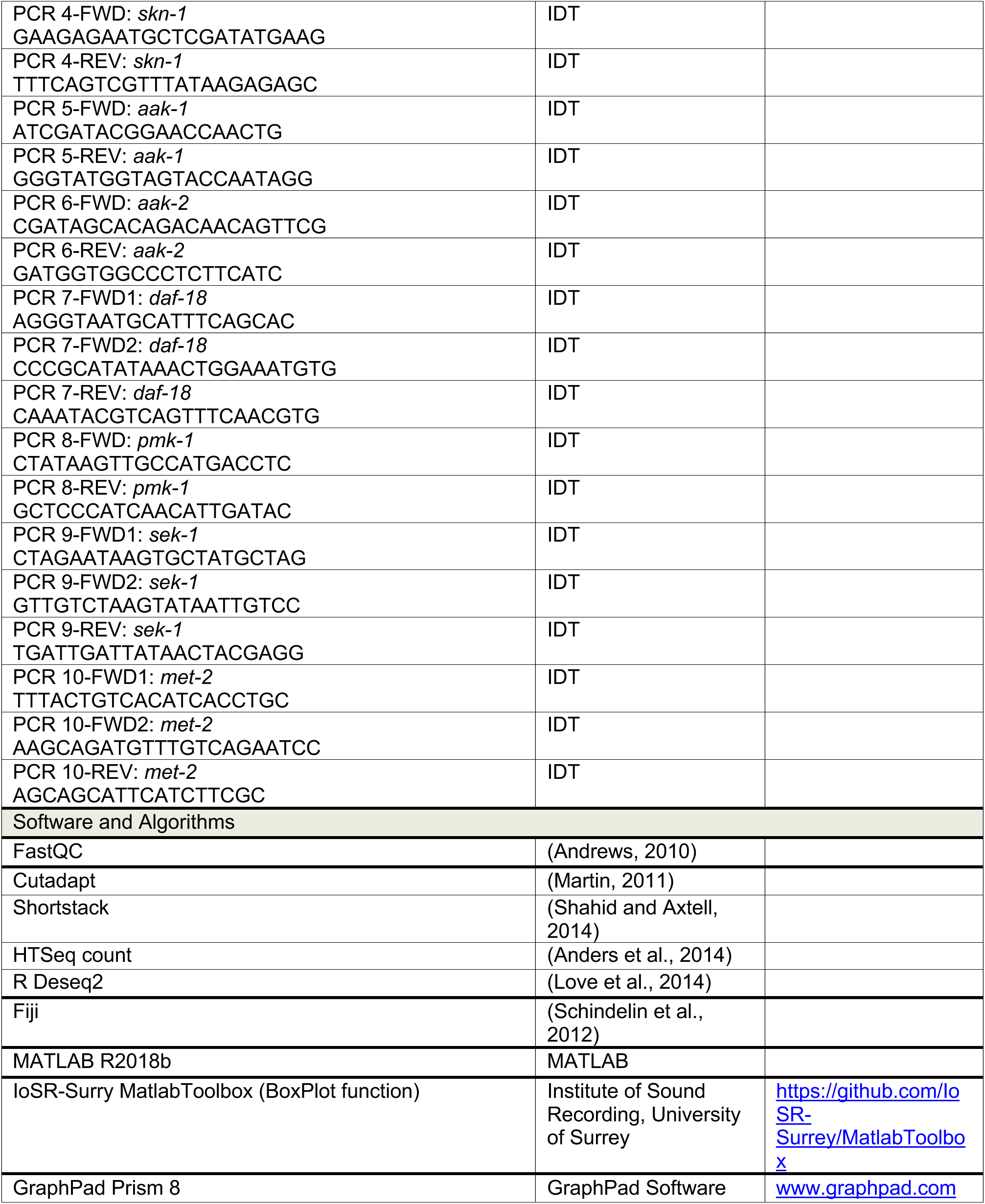

